# Conserved amino acids residing outside the voltage field can shift the voltage sensitivity and increase the signal size of *Ciona* based GEVIs

**DOI:** 10.1101/2020.08.27.269761

**Authors:** Masoud Sepehri Rad, Lawrence B. Cohen, Bradley J. Baker

## Abstract

To identify potential regions of the voltage-sensing domain that could shift the voltage sensitivity of *Ciona intestinalis* based Genetically Encoded Voltage Indicators (GEVIs), we aligned 183 voltage-gated sodium channels from different organisms. Conserved polar residues were identified at multiple transmembrane loop junctions in the voltage sensing domain. Similar conservation of polar amino acids was found in the voltage sensing domain of the voltage-sensing phosphatase gene family. These conserved residues were mutated to nonpolar or oppositely charged amino acids in a GEVI that utilizes the voltage sensing domain of the voltage sensing phosphatase from *Ciona* fused to the fluorescent protein, super ecliptic pHlorin (A227D). Different mutations shifted the voltage sensitivity in a more positive or a more negative direction. Double mutants were then created by selecting constructs that shifted the optical signal to more negative potentials resulting in an improved signal in the physiological voltage range. The combination of two mutations, Y172A (intracellular loop between transmembrane segments S2 and S3) and D204K (extracellular loop between transmembrane segments S3 and S4) showed the increased the signal size from 2.5% to 13.8% for a 100 mV depolarization. Introduction of these mutations into previously developed GEVIs improved the dynamic range to 40% ΔF/F/100 mV.

## INTRODUCTION

Genetically encoded voltage indicators (GEVIs) are potentially powerful tools for monitoring electrical activity in the brain. Fusing a green fluorescent protein (GFP) to the voltage-activated Shaker potassium channel Siegel and Isacoff developed the first archetype, Flash [1]. Kinetics, brightness and the signal size are some of the important properties of a GEVI [2, 3]. Improving properties of the voltage sensors is necessary for increasing the utility of GEVIs for imaging fast electrical activities in neural tissue and *in vivo* [4, 5]. GEVIs with fast kinetics and large dynamic signals are needed to follow the voltage transients of neurons. In recent years, several attempts have been made to improve GEVI’s properties [6-15].

The GEVI, ArcLight, yields a large change in fluorescence in response to changes in membrane potentials [16]. ArcLight can give up to a 40% ΔF/F for **a** 100 mV depolarization of the plasma membrane in HEK 293 cells [17]. However, its fast time constant is ∼ 10 ms. Since action potentials are 1-2 ms, ArcLight will only reach ∼10% of its maximal signal by the time the spike has subsided. The GEVI, Bongwoori, is an ArcLight-derived construct with improved speed (tau – 8 ms) and a shifted voltage sensitivity towards positive potentials [9]. By focusing on the voltage sensing domain (VSD) of the *Ciona* VSP, conserved polar amino acids were identified in transmembrane segments that affect the voltage sensitivity of the probe. Introducing mutations in these locations affected the voltage range, the speed of optical response, and the signal size of the GEVI. The resulting probe, Bongwoori, had sufficient speed and sensitivity to resolve action potentials in a hippocampal neuron firing at 60 Hz. Further improvement of the Bongwoori family of GEVIs was achieved by altering the charge composition of the region linking the fluorescent protein (FP) to the VSD domain [8]. Introduction of positively charged residues in the linker region improved the signal size of Bongwoori yielding fluorescent signals as high as 20% ΔF/F during action potentials.

The Fluorescence versus voltage relationship is an important factor for a GEVI. For example, the GEVI CC1 based on the wildtype sequence of the *Ciona* phosphatase gives a modest signal for a 200 mV depolarization step but gives only a small optical signal for a 100 mV step [9]. Shifting the voltage sensitivity of CC1 to more negative potentials may improve the signal size in the physiological voltage range. Here, we show that the mutagenesis of conserved polar amino acids in intra/extracellular loops in the VSD can affect the voltage range of the *Ciona* based GEVIs even though they reside outside of the voltage field. Making double mutant constructs, we have successfully shifted the voltage sensitivity of CC1 to more negative potentials and improved its signal size for physiologically relevant voltage ranges.

## RESULTS

### CC1 mutations

The GEVI, CC1, contains the wild type VSD sequences of the *Ciona intestinalis* voltage sensing phosphatase gene with the FP, SuperEclipticpHlorin A227D, fused to the carboxy-terminus of the protein. Introduction of mutations in the α-helix/loop junctions in the VSD of CC1 yielded probes with shifted voltage sensitivity in a more positive or a more negative direction. To identify other regions of the VSD which contribute to determining the voltage sensitivity, 183 voltage-gated sodium channels (Nav) from different organisms were aligned. Nav channels were chosen since they have four VSDs in each protein. Conserved polar residues were detected at multiple transmembrane/loop junctions (Fig. 1). These conserved residues were mutated to nonpolar or oppositely charged amino acids.

**Fig. 1.**
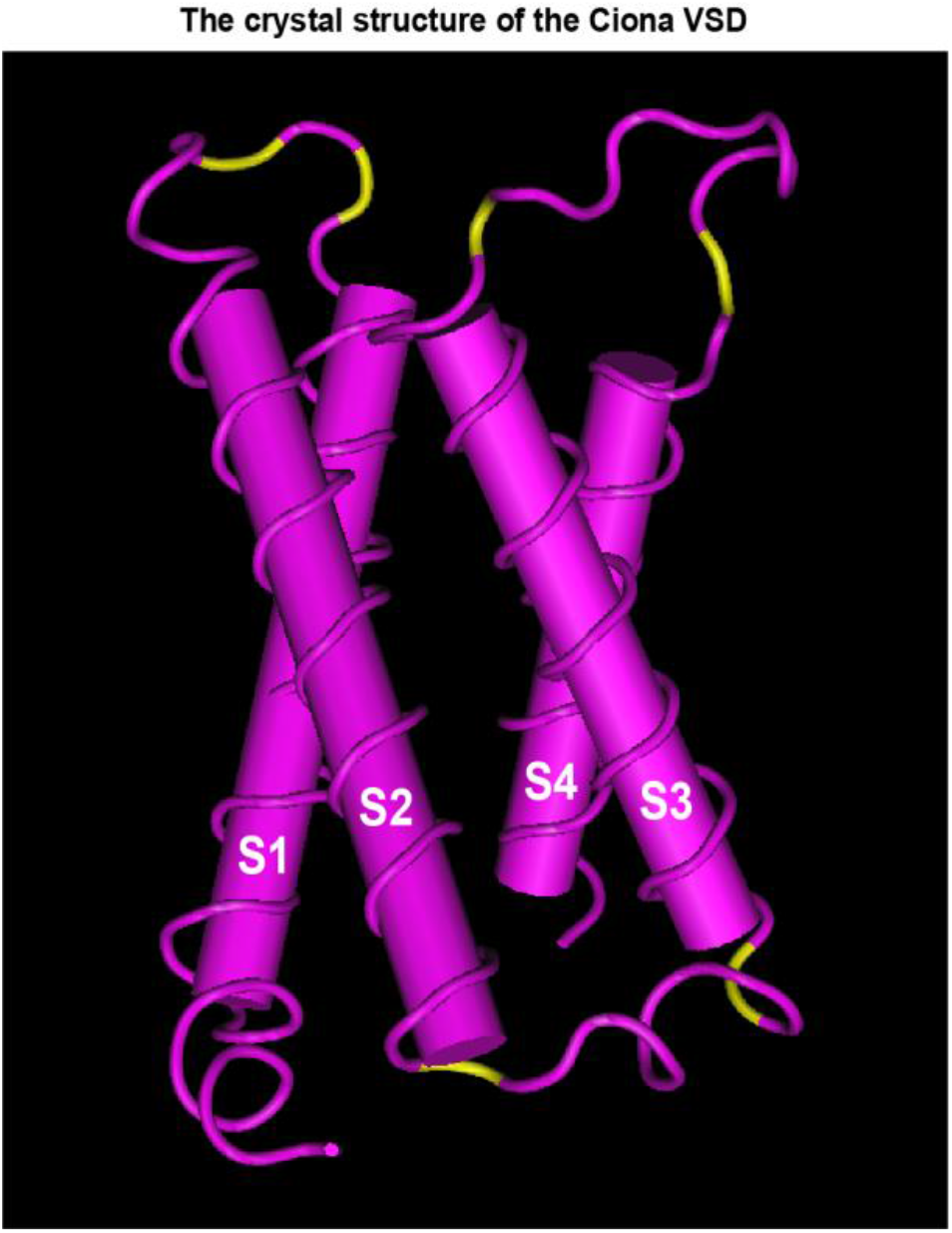
Crystal structure of Ciona VSP [18]. Yellow regions depict amino acid positions that were mutated in this study.

#### S1-S2 loop mutants

Figure 2 compares the traces of the S1-S2 loop mutants with that of the CC1 construct. Except I140S, none of the S1-S2 loop mutations shifted the voltage sensitivity significantly. I140S shifted the voltage at half-maximum signal from 70 mV to 48 mV (Fig. 2B). While I140A, E and K142A, S decreased the signal size for a 100 mV depolarization, I140S increased the signal size from 2.5% to 3.5% (Fig. 2A).

**Fig. 2.**
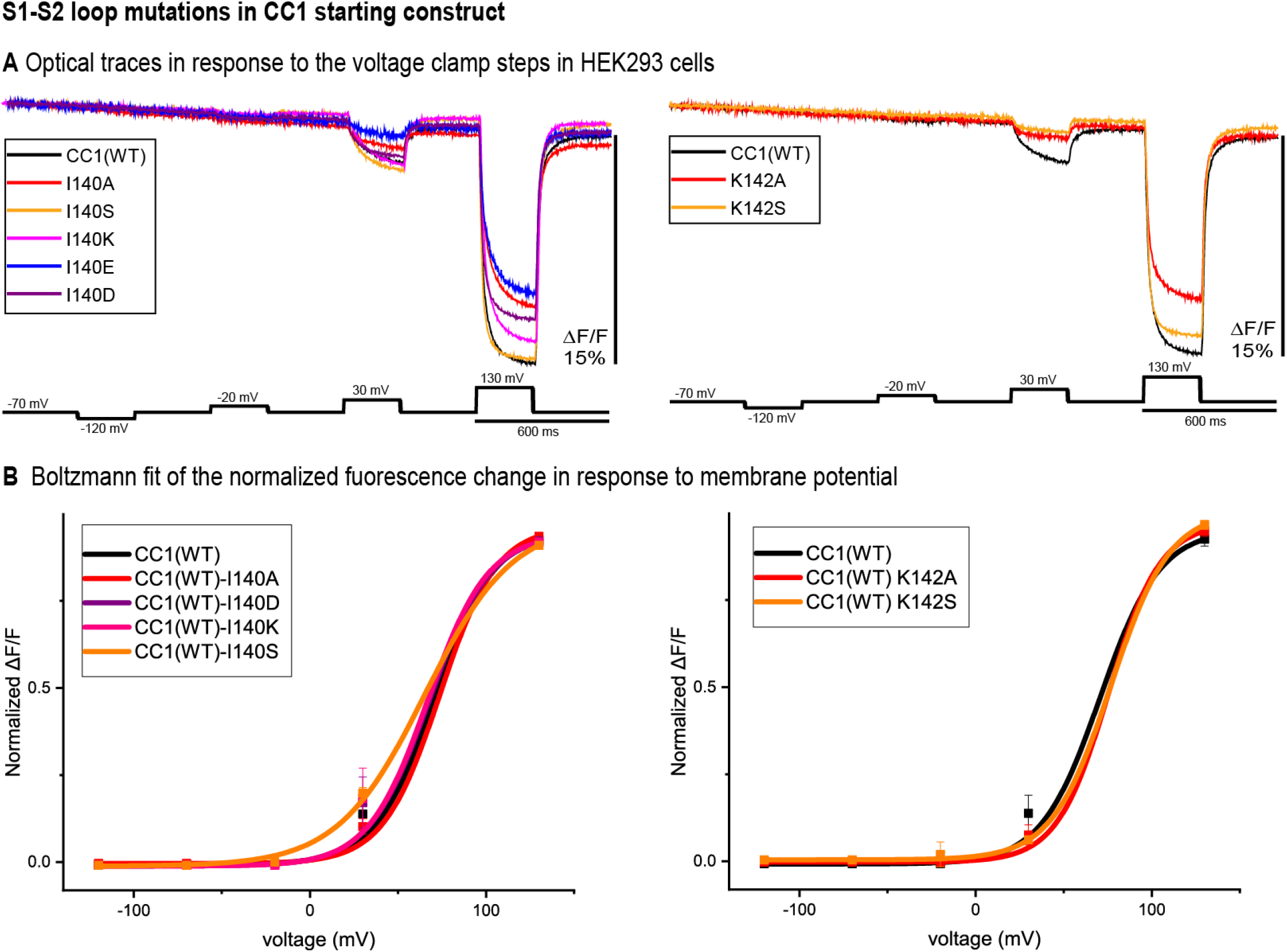
S1-S2 loop mutations in the CC1 starting construct. **A.** The traces are representative fluorescent signals from voltage-clamped HEK293 cells (average of 16 trials for each cell) expressing CC1(WT) or CC1(WT) with an S1-S2 loop mutant. Each trace is the average of three cells. The holding potential was −70mV. The pulse protocol is given in the black trace and was identical for all constructs. No temporal filtering was used for the traces. Images were recorded at a frame rate of 500 fps. **B.** Boltzmann fit of the normalized fluorescence change in response to membrane potential (n=3 for all constructs).

#### S2-S3 loop mutants

Figure 3 indicates the optical signal traces of the intracellular S2-S3 loop mutants in comparison to the CC1 starting construct. While Y172D did not change the signal size for a 100 mV depolarization, Y172A, S, K, and R did increase the signal size compared to the CC1. Y172R and Y172K increased the signal size from 2.5% to 4.8% and 5% in this location respectively. Most of the mutations at this location did not change the signal size dramatically. N180D had a deleterious effect such that it decreased the signal size from 2.5% to 0.8% for **a** 100mV step (Fig. 3A). We used the Boltzmann fit to the CC1 construct as a reference point for the mutations to the loop junctions (Fig. 3B). All of the Y172 mutations tested shifted the voltage response to more negative potentials, with Y172A having the largest effect. This single mutation shifted the voltage at half maximum from 70 mV to 45 mV. The mutations in the other side of the S2-S3 loop junction (N180) did not change the voltage sensitivity significantly (Fig. 3B).

**Fig. 3.**
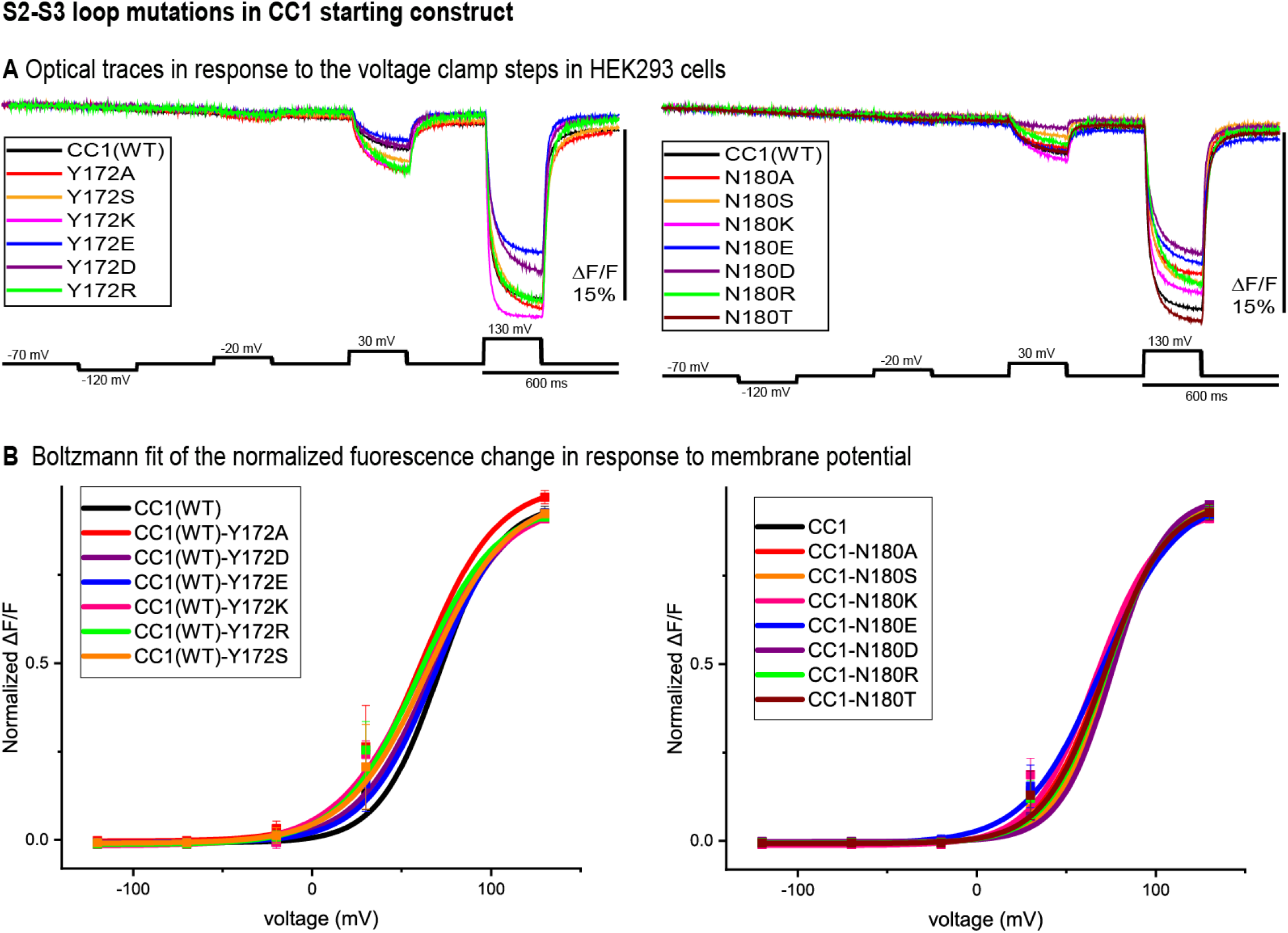
S2-S3 loop mutations in the CC1 starting construct. **A.** Representative fluorescent traces of HEK cells expressing the CC1(WT) or CC1(WT) with an S2-S3 loop mutant in the whole cell patch clamp configuration. The cells were subjected to the voltage command pulses in black. Each trace is from average of three cells and the results from each cell is the average of 16 trials. The traces are fluorescent optical signals from plasma membrane without temporal filtering. Images were recorded at a frame rate of 500 fps. **B.** Boltzmann fit of the normalized fluorescence change for the CC1 construct and its derivative probes with single mutation in the S2-S3 loop (n=3 for all constructs).

#### S3-S4 loop mutants

Figure 4, represents the optical signal traces from the extracellular S3-S4 loop mutants. D204S, K, E, R, T, and Y have a larger signal size for 100mv depolarizations compared to the CC1 signal; D204K has the largest effect increasing the signal size from 2.5% to 8.1%. D213A, S, K, T, Y and D204A have a smaller signal size for 100mv depolarizations (Fig. 4A). Two CC1 derived probes with a single mutation in this location (CC1-D213E and CC1-D213R) did not express in HEK293 cells (data not shown). While the D204A mutation did not shift voltage vs fluorescence, all of the other mutations tested in this location shifted the voltage response of the CC1 probe to more negative potentials. D204K had the most pronounced effect shifting the V1/2 to from +70 mV to +17 mV. While the D204 mutations exhibited a shifted voltage sensitivity, the D213 mutations did not. (Fig. 4B).

**Fig. 4.**
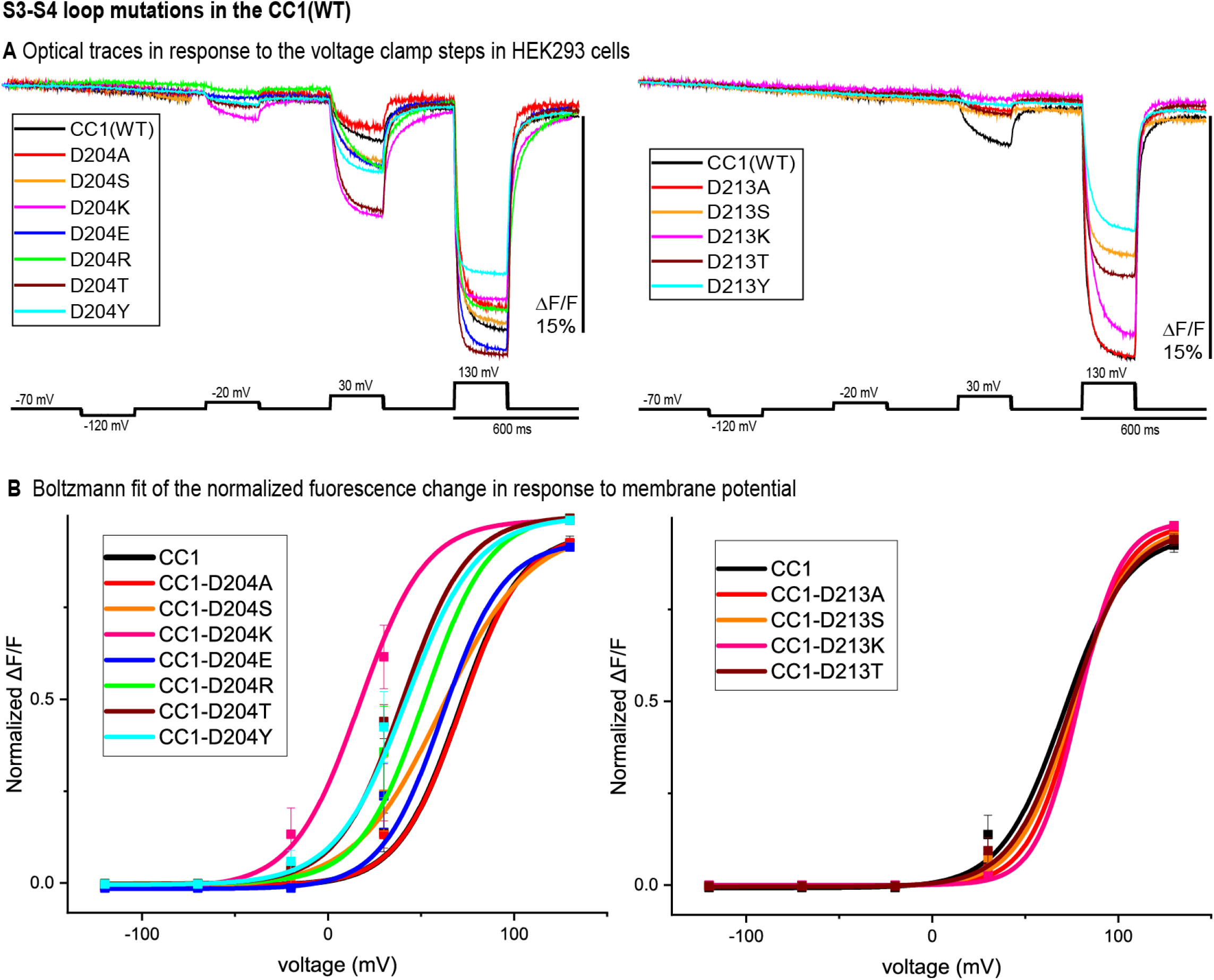
S3-S4 loop mutations in the CC1 starting construct. **A.** HEK 293 cells expressing the CC1(WT) or CC1(WT) with an S3-S4 loop mutant were voltage-clamped. The traces are fluorescent optical signals (average of three cells) from plasma membrane without temporal filtering. Each cell is the average of 16 trials. Images were recorded at a frame rate of 500 fps. **B.** The voltage-fluorescence curve of the normalized optical signal. for the CC1 construct and its derivative probes with single mutation at S3-S4 loop (n=3 for all constructs).

#### Double mutants

One of the goals in the development of a protein-based voltage probe is increasing the optical signal size. The CC1 starting construct has a voltage response shifted to more positive potentials compared to the physiological voltage range (−70 mV to +30 mV). We tried to further increase the signal size for a 100 mv depolarization by shifting the voltage sensitivity to even more negative potentials using double mutations. Comparing optical traces of the loop mutants with that of the starting CC1 construct we selected constructs that showed relatively large voltage shift**s** to more negative potentials and made four new probes. The voltage response of these double mutant constructs are shifted to even more negative potentials and as a result they give a larger signal compared to CC1 for a 100 mv depolarization (Figure. 5A). Y172A/D204K had the largest effect (Figure. 5B). This construct increased the signal size for 100mv depolarizations compared to the CC1 optical signal from 2.5% to 13.8% for a 100 mv depolarization.

**Fig. 5.**
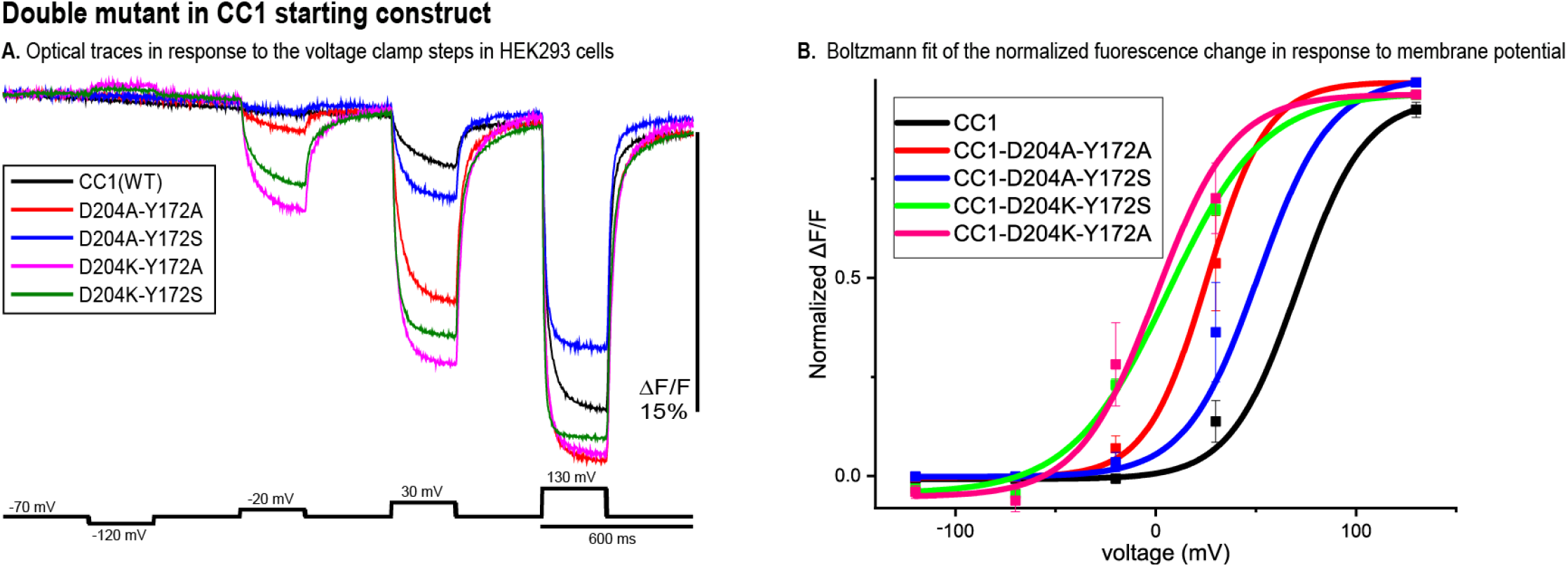
Double mutant in the CC1 starting construct. **A.** Fluorescence response of HEK293 cells expressing CC1(WT) or CC1(WT) with double mutations in the loops to depolarizing and hyperpolarizing voltage steps. The traces are the average of three cells (each cell is the average of 16 trials) without temporal filtering. Images were recorded at a frame rate of 500 fps. **B.** The optical signal was normalized and fitted to a Boltzmann equation. All double mutant combinations shifted the voltage response to more negative potentials (n=3 for all constructs).

Introducing the double mutant Y172A/D204K as well as the D164N into Pos6 [8], we made a new GEVI (named Plos6-v2). Plos6-v2 has a large signal size (∼40% for 100 mV depolarizations) (Figure 6). The average signal size from three cells for each of the four double mutants in Figure 5 is D204K-Y172A, 34.1%; D204k-Y172K, 33.2%; D204K Y172R, 29.0%; and D204K Y172S, 35.8%. These values were obtained by averaging the pixels around the edge of each of the HEK293 cells. If the signal from the whole cell is used, the fractional fluorescence change was smaller by ∼4% because of the fluorescence from internal probe expression which, for these GEVIs, we presume does not change in response to the change in membrane potential.

**Fig.ure 6.**
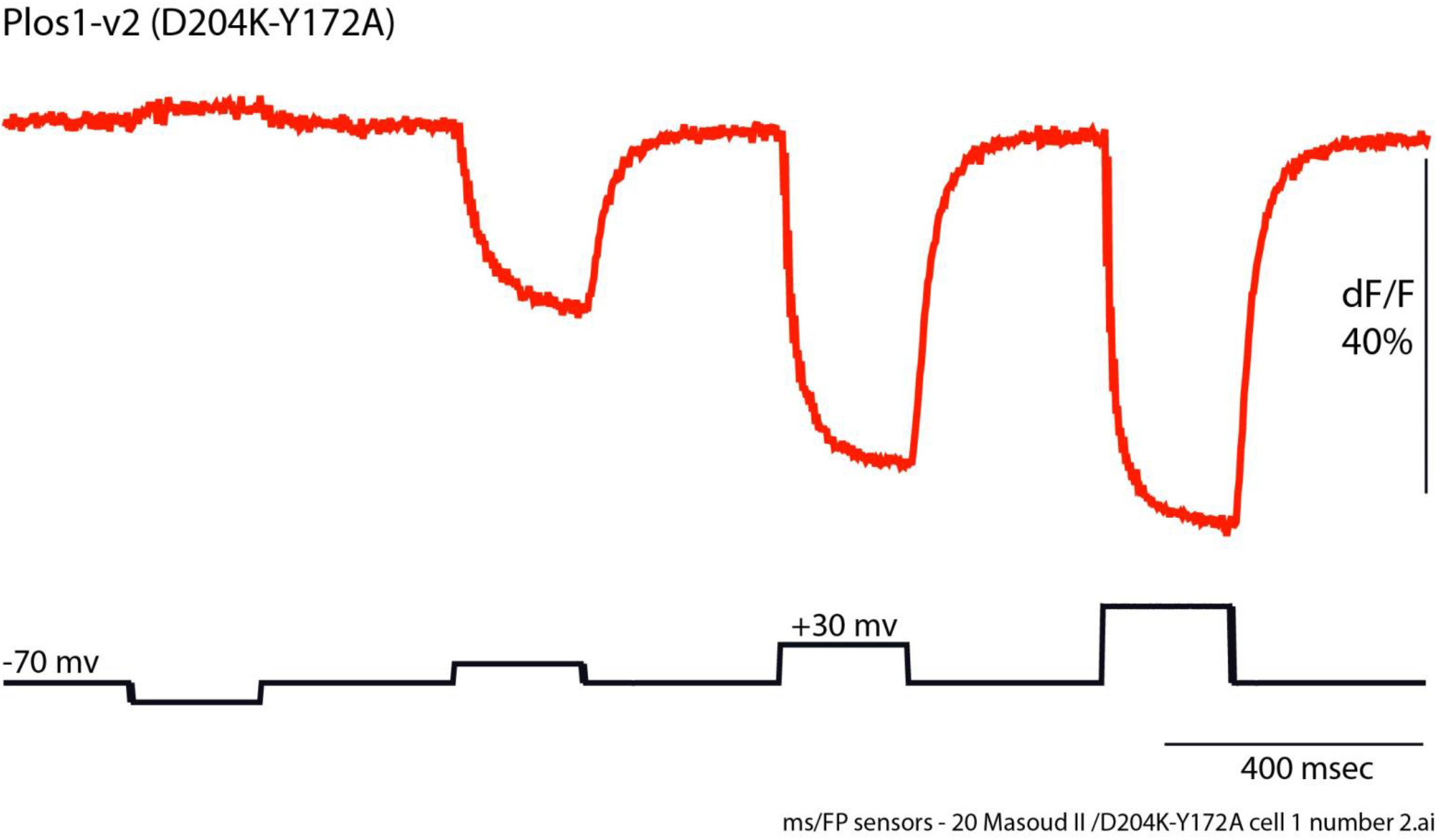
The relatively large fluorescence response of Plos6-v2 in a HEK293 cell. The fractional fluorescence change for a 100 mv depolarization was ∼40%. The fluorescence trace is from a single trial. The NeuroCCD camera frame rate was 500 Hz; no temporal filtering was used.

## DISCUSSION

Piao et al. (2015) showed that mutagenesis of conserved polar amino acids in the transmembrane segments of the VSD affects the voltage sensitivity of *Ciona* based GEVIs In this report, we have shown that the conserved polar residues in the transmembrane loop junctions can also affect the characteristics of the voltage-dependent optical signal. We identified several mutations in the α-helix/loop junctions in the VSD that shift the voltage sensitivity of the GEVI. In the S1-S2 loop re**g**ion I140A,E and K142A,S mutations shifted the voltage sensitivity to more positive voltages. While, I140S,K and D did not change the voltage sensitivity dramatically. In the intracellular S2-S3 loop, Y172A,S,K and R shifted the voltage response to more negative and N180S,R,T shifted the voltage response to more positive potentials. Mutagenesis of D204S, K, E, R, T, and Y in S3-S4 loop shifted the voltage response to more negative potentials, while D204A and D213A, S, K, T, and Y mutations shifted the voltage response of the optical signal to more positive potentials. The mechanism by which the loop mutations shift the voltage sensitivity is not well understood.

Selecting constructs that showed the largest voltage shift to more negative potentials, we made four double mutant probes. The voltage response of these new GEVIs are shifted to even more negative potentials and as a result they give a larger signal compared to CC1 for a 100 mv depolarization (Figure. 5A). Y172A/D204K showed the biggest effect, it increased the signal size from 2.5% to 13.8%. Introducing positively charged residues in the linker region, an ArcLight derived probe, Pos6 was developed. Here, we introduced D164N/Y172A/D204K mutations in Pos6 in an attempt to increase its optical signal size even further. The resulting construct gives nearly a 40% ΔF/F optical signal for a 100 mV depolarization which is about a 4-fold improvement over the original CC1-Pos6 construct.

## Materials and Methods

### Plasmid DNA designs and construction

The CC1(WT) construct was described in Piao et al., 2015. Double mutant probes (CC1(WT)-D204K-Y172A and CC1(WT)-D204K-Y172S) were made by a two-step PCR process using CC1(WT)-Y172A and CC1(WT)-Y172S as template DNA respectively. Primers used for amplification of the S1, S2 and S3 transmembrane domains with a single mutation (D204K) in first step PCR reaction were: LC226: 5-ATA CGA CTC ACT ATA GGG-3 and LC230: 5- accGTATTCCTTTAACACAGT -3. Primers used for amplification of S4 transmembrane domain and florescence protein in first step PCR reaction were: LC229: 5- ACT GTG TTA AAG GAA TAC ggt -3 and LC193: 5- GCGATATCTTCTTTTGTTaaaa -3. In the second step PCR, we used primers LC226 and LC193 and combined the first step PCR products. The second step PCR product then was digested with restriction enzymes Nhe1 and Kpn1 and inserted into the corresponding sites of the CC1(WT) construct. Similarly, we used CC1(WT)-Y172A and CC1(WT)-Y172S as template DNA respectively to generate the double mutant probes (CC1(WT)-D204A-Y172A and CC1(WT)-D204A-Y172S). For amplification of the S1, S2 and S3 transmembrane domains with a single mutation (D204A) in first step PCR reaction these primers were used: LC226: 5-ATA CGA CTC ACT ATA GGG-3 and LC232: 5- accGTATTCGGCTAACACAGT -3. Primers used for amplification of S4 transmembrane domain and florescence protein in first step PCR reaction were: LC231: 5- ACT GTG TTA GCC GAA TAC ggt -3 and LC193: 5- GCGATATCTTCTTTTGTTaaaa -3. We then used primers LC226 and LC193 to combine the first step PCR products. Using enzymes Nhe1 and Kpn1, the second step PCR product then was digested and inserted into the corresponding sites of the CC1(WT) construct. All DNA constructs were confirmed by DNA sequencing (Cosmogenetech, Republic of Korea).

### Cell culture

HEK293 cells were maintained in DMEM (High Glucose DMEM; Gibco) supplemented with 10% (v/v) fetal bovine serum (FBS; Invitrogen). HEK293 cells were seeded on to #0 coverslips coated with poly-L-lysine (Sigma) in a 24-well culture dish and kept in an incubator at 37°C under air with 5% CO2. Transfection was performed by using Lipofectamine 2000 (Invitrogen) following the manufacturer’s instructions.

### Patch clamp

Electrophysiology recordings were performed at 33 °C and the chamber was perfused with a bath solution containing 150 mM NaCl, 4 mM KCl, 2 mM CaCl2, 1 mM MgCl2, 5 mM D-Glucose, and 5 mM HEPES (pH 7.4). Glass patch pipettes (capillary tubing with 1.5/0.84 mm; World Precision Instruments) were pulled by a P-97 micropipette puller (Sutter Instruments, USA) to make patch pipettes with 3–5 MΩ resistance when filled with internal solution containing (in mM) 120 K-aspartate, 4 NaCl, 4 MgCl2, 1 CaCl2, 10 EGTA, 3 Na2ATP, and 5 HEPES, pH 7.2. Using a Patch Clamp EPC10 amplifier (HEKA) with a holding potential of -70 mV, whole-cell voltage clamp was done in transfected HEK293 cells.

### Wide-field imaging

We used an inverted microscope (IX71; Olympus, Japan) equipped with a 60X oil-immersion lens with 1.35-numerical aperture (NA) for imaging the whole-cell patch clamped cells. Illumination light was provided by a 75 W xenon arc lamp (Cairn Research). The excitation filter for all constructs was 472/30, the emission filter was 496/LP and the dichroic was 495 (Semrock, NY). The fluorescence image was demagnified by an Optem zoom system, A45699 (Qioptiq LINOS) and the sample imaged onto a NeuroCCD-SM camera with 80×80 pixels controlled by NeuroPlex software (RedShirtImaging, GA). The images were recorded at a frame rate of 500 fps.

### Optical signal analysis

Acquired images from patch clamp fluorometry were analyzed using Neuroplex software (RedshirtImaging, USA), Excel (Microsoft, USA), and Origin8.6 (Origin Labs, USA). The resulting traces from whole cell voltage clamp experiments of HEK 293 cells were averaged for 16 trials. To calculate the % ΔF/F, we first subtracted the dark image from all frames, then the average of a region of interest in each frame during a voltage step (F) was subtracted from the average of the region taken from ten frames prior to the event of interest (F0), and finally this value was divided by F0 by the following formula: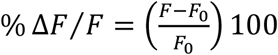. The optical traces were imported into Origin 8.6 (OriginLab, MA) for the analysis of response time constants, *τ*_1_ and *τ*_2_. The optical traces then were fitted with either a single exponential equation 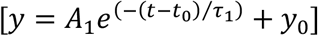 or a double exponential equation 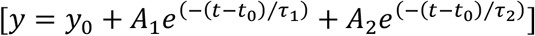 where *A*_1_ and *A*_2_ are amplitudes, and t is time in milliseconds. To compare the optical responses that were better fitted to a double exponential decay to those that could only be fit by a single-exponential decay, a weighted tau was calculated by the following formula: [*τ*_*weighted*_ = *τ*_1_(*A*_1_/(*A*_1_ + *A*_2_)) + *τ*_2_(*A*_2_/(*A*_1_ + *A*_2_))]. ΔF/F values for all of the tested constructs were plotted in OriginPro 2019 (OriginLab, USA) and fitted to a Boltzmann function (signal size was normalized from zero, minimum, to one, maximum) to acquire voltage sensitivities as previously described [9].

## Acknowledgements

This work was supported by World Class Institute (WCI) Program of the National Research Foundation of Korea (NRF) funded by the Ministry of Education, Science and Technology of Korea (MEST) (NRF Grant Number: WCI 2009-003), the Korea Institute of Science and Technology (KIST) grants 2E26190 and 2E26170 and NIH grants DC005259 and U01NS099691. The content is solely the responsibility of the authors and does not necessarily represent the official views of the National Institutes of Health.

